# Synaptic cleft geometry modulates NMDAR opening probability by tuning neurotransmitter residence time

**DOI:** 10.1101/2024.10.30.621190

**Authors:** M. Hernández Mesa, K. J. McCabe, P. Rangamani

## Abstract

1

Synaptic morphology plays a critical role in modulating the dynamics of neurotransmitter diffusion and receptor activation in interneuron communication. In this study, we investigated how variations in synaptic geometry, including curvature of the synaptic cleft, distance between the presynaptic and postsynaptic membranes, and the surface area-to-volume ratio of the cleft, influence glutamate diffusion and N-Methyl-D-Aspartate receptor (NMDAR) opening probabilities. We developed a stochastic model for receptor activation using reconstructions from realistic synaptic geometries. Our simulations revealed a substantial variability in NMDAR activation, highlighting the significant impact of synaptic structure on receptor dynamics. By exploring the interplay between curvature and surface area-to-volume ratio, we found that increasing the curvature of the synaptic membranes could compensate for reduced NMDAR activation when the synaptic cleft distance was large. We also found that non-parallel membrane configurations, particularly convex presynapses or concave postsynaptic densities (PSDs), maximize NMDAR activation via increased surface area-to-volume ratio, leading to prolonged glutamate residence and reduced leakage. Finally, introducing NMDAR clustering within the PSD significantly enhanced receptor activation across different geometric conditions and mitigated the effects of synaptic morphology on NMDAR opening probabilities. Our findings underscore the complex interplay between synaptic geometry and receptor dynamics, providing insights into how structural modifications can influence synaptic efficacy and plasticity.

**Significance statement:** This study demonstrates that synaptic morphology profoundly shapes neurotransmitter diffusion and NMDA receptor activation, directly impacting synaptic efficacy. Our model shows that factors like synaptic cleft curvature, membrane spacing, and surface area-to-volume ratio significantly influence receptor dynamics. Given the dynamic nature of dendritic spines, which change shape and size during synaptic plasticity, our findings illustrate how purely morphological changes in cleft structure can modulate interneuronal communication and signal strength.

## 3 Introduction

Communication between neurons occurs at specialized contact points known as synapses. A synapse consists of a presynaptic terminal and a postsynaptic membrane located on two different neurons. The gap between these neurons, known as the synaptic cleft, facilitates the release of neurotransmitters from the presynaptic terminal to activate receptors on the postsynaptic membrane (*1*). The distance of the synaptic cleft distance has been measured to be approximately 12 to 50 nm (*2–4*). During excitatory synaptic transmission in glutamatergic neurons, the presynaptic terminal releases the neurotranmsitter glutamate into the synaptic cleft, allowing for interneuron communication (*5*). The postsynaptic density (PSD) is a specialized protein-rich region of the postsynaptic membrane and contains glutamate receptors. Among the receptors within the PSD, the glutamatergic receptor N-methyl-D-aspartate (NMDAR) is particularly important for synaptic plasticity. NMDAR activation requires simultaneous glutamate binding and membrane depolarization (*6*). Under these combined conditions, NMDARs allow the entry of Na^+^ and calcium ions (Ca^2+^) into the spine (*5*). Influx of Ca^2+^ through NMDARs initiates a cascade of synaptic signaling pathways that are relevant in the transduction of neural signals (see overviews in (*7–10*)). Spines and their glutamatergic receptors, particularly NMDARs, play a central role in synaptic plasticity, regulating learning and memory formation.

The geometry of the synaptic cleft can modulate the diffusion of neurotransmitters released from the presynaptic terminal, impacting the probability of NMDAR activation. Synaptic morphology has been widely studied to understand how it evolves during synaptic plasticity and affects on downstream signal transduction (*2, 11*). Changes in synaptic morphology, such as alterations in the spine size and the surface area of the postsynaptic density (PSD), influence the spatial arrangement of NMDARs within the synaptic cleft. Experimental measurements have shown a wide range of PSD area, varying from 0.006 *μ*m^2^ to 0.334 *μ*m^2^ (*4, 12–15*). Synaptic curvature is often associated with the efficiency of vesicle docking and neurotransmitter release (*11, 16*). For instance, Marrone et al. (*13*) demonstrated that synaptic membranes displaying higher curvature had a greater number of docked vesicles compared to those with flatter profiles. These observations suggest a connection between membrane curvature, synaptic plasticity, and synaptic transmission efficiency. It is widely recognized that cells typically adapt their shapes according to their specific cellular processes and functions (*17–19*). Notably, curvature is a physical phenomenon that is relevant in biological systems across multiple scales, from proteins (*20, 21*) to sub-cellular membrane shape, organelle shape, and tissue level structures (*22*). Previous, we and others have shown that spine shape can affect calcium signaling and synaptic plasticity in dendritic spines (*7, 8, 23–25*). Building on our previous findings (*23*), which revealed significant changes in NMDAR opening probability across three different synaptic cleft geometries, we decided to carry out a larger study on different realistic synapses. With this in mind, our objective is to advance our understanding of how the shape of the synaptic cleft influences neural communication.

In what follows, we refer to concave synapses when the presynaptic membrane extends into the postsynaptic density, while the term convex synapses describes synapses where the postsynaptic density protrudes into the presynaptic membrane, as in (*2, 11*). A graphical description of this terminology is given in Figure 1. The curvature of the PSD is altered during synaptic potentiation (*11*). In an early study, Markus and Petit observed a shift from a balance between concave and convex synapses in the early stages of development, to a predominance in convex synapses in mature adult rats (*16*). Several studies have reported an increase in the occurrence of concave synapses 24 hours after long term potentiation (LTP) induction (*11, 26–28*). Similar results were also reported within minutes after LTP induction (*14, 29*). Medvedev et al. (*30*) observed that synapses in mushroom spines became less concave (increasingly flatter) after LTP, while thin spines showed very pronounced changes in PSD curvature, transforming concave synapses into convex ones (*30*). These changes align with the observations of Markus and Petit who observed more convex spines in mature adult rats (*16*). Such variations in synaptic morphology underscore the importance of spine shape in modulating synaptic function.

**Figure 1:**
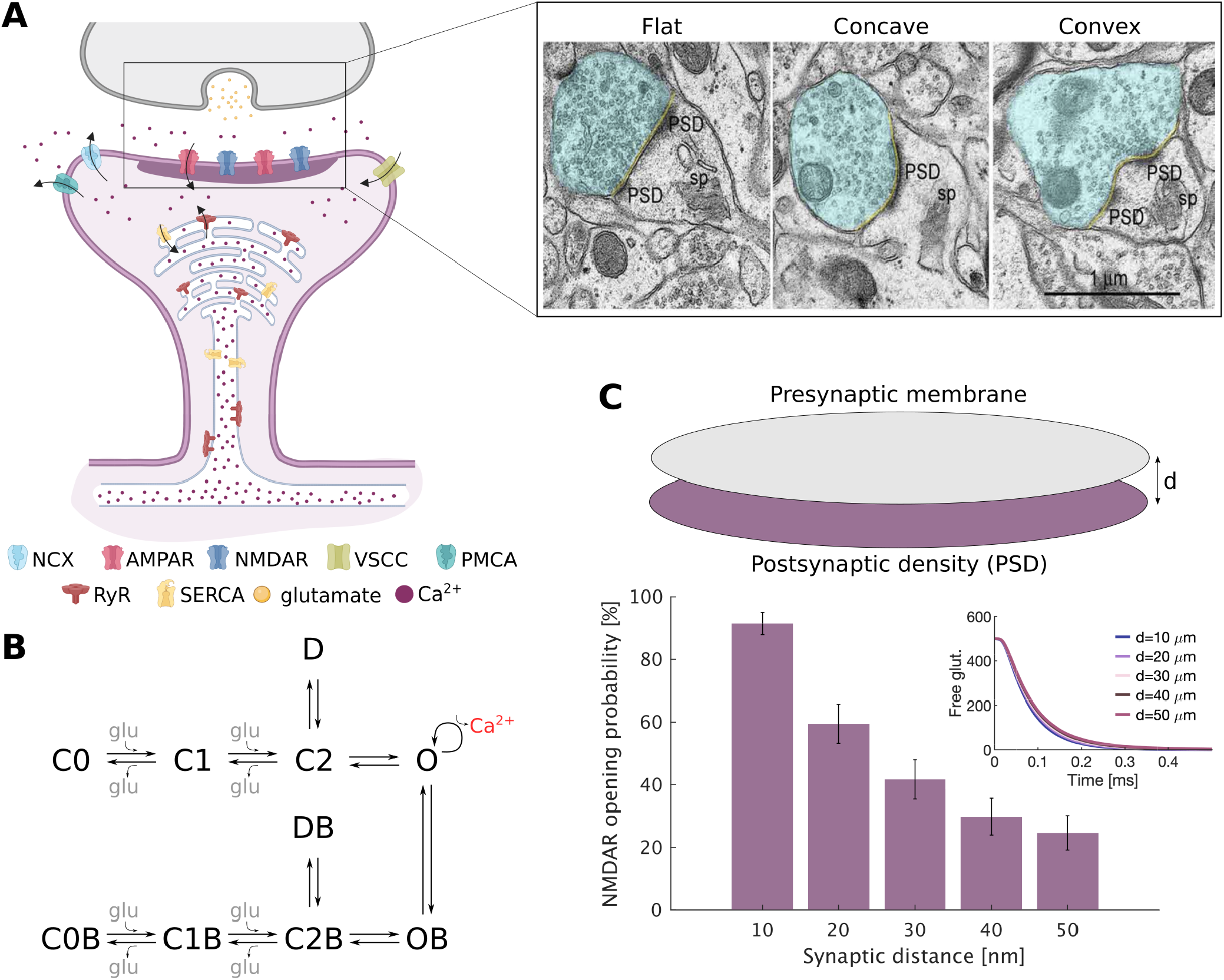
Overview of the geometric variables in the synaptic cleft and effect of synaptic cleft distance on NMDAR probability. **A**: Schematic of the signal transmission in a synapse. Parts of this panel were generated using *Biorender*.*com*. This geometry was taken from the mesh generated by Lee et al. (*31*) from images taken by Wu et al. (*32*). The inset to the right is an image taken from (*30*) showing different synaptic curvatures measured in adult rats using electron microscopy. *Permission pending for image use from Medvedev et al.* (30). **B**: Schematic of NMDAR reaction rate model from Vargas-Caballero and Robinson (*33*). **C**: Simulation show the effect of synaptic cleft distance on NMDAR opening probability and free glutamate within the synapse.

We build on the prior work of Rusakov et al. (*34*) where they explored how glutamate diffuses within the synaptic cleft and how the release distance from the cleft affects NMDA and *α* -amino-3-hydroxy-5-methyl-ioxazolepropionate (AMPA) receptor activation. Furthermore, they showed how sensitive NMDARs are to the duration of glutamate diffusion at micromolar levels (*34*). Here, we investigated how cleft distance, and cleft morphology and curvature affects NMDAR activation both in realistic and idealized geometries using stochastic modeling. In this work, we developed particle-based stochastic spatial models to investigate how synaptic cleft morphology influences NMDAR activation. Suprisingly, we found that realistic geometries exhibit significant variability in NMDAR receptor activation.

To understand the sources of this variability, we focused on key geometric variables such as synaptic membrane curvature, distance between the membranes, and surface-to-volume ratio to identify how these features can affect the residence time of glutamate and NMDAR activation. Using idealized geometries, we found that increasing synaptic membrane curvatures could increase NMDAR activation probability. We also showed that maintaining a constant surface area-to-volume ratio while varying synaptic cleft distances resulted in higher NMDAR opening probabilities at intermediate distances but decreased activation at short and long distances. Finally, we demonstrate that receptor clustering could have an important physiological role in overcoming such variation. Thus, our study gives insights into how synaptic cleft morphology affects neurotransmitter residence time and NMDAR activation.

## 4 Model development

In this work, we use a particle-based stochastic modeling approach to study the effects of geometrical features on particle diffusion in enclosed volumes delimited by membranes, such as a synaptic cleft. We integrate both idealized and realistic geometries reconstructed from experimental images. We outline the main steps in our model below.

### 4.1. Geometries

#### 4.1.1. Realistic geometries

We adopted the synaptic structures segmented and meshed by Lee et al. (*31*). These synapses were reconstructed from neurons in the mouse cerebral cortex, imaged using focused ion beam scanning electron microscopy, as detailed in the work by Wu et al. (*32*). With particular interest in the PSD, we directed our attention to the PSD detailing within the reconstructions provided by Lee et al. (*31*). A subset of the reconstructed spines are depicted in Figure 2A. To simulate a complete synaptic structure, and considering that only the PSD components were reconstructed, our model presumes an identical membrane geometry for both the PSD and the presynaptic terminal. We simulated this placing an identical membrane in parallel with the PSD at a gap of 17.32 nm, which is corroborated by empirical measurements reported in the literature (*2, 3*). Given our modeling assumption that the presynaptic terminal is aligned parallel to the PSD and that the two membranes are identical structures, we ensure that the separation between any corresponding pair of points across the membranes is consistently maintained at 17.32 nm. The location of glutamate release was chosen by calculating the geometrical center of the presynaptic terminal. To ensure that the neurotransmitter was optimally released between the presynaptic terminal and the PSD, we identified the mesh point on the presynaptic terminal closest to this geometric center. The precise release point of the neurotransmitter was then positioned at 0.01 nm from the identified mesh point. 500 glutamate molecules were released at this site.

**Figure 2:**
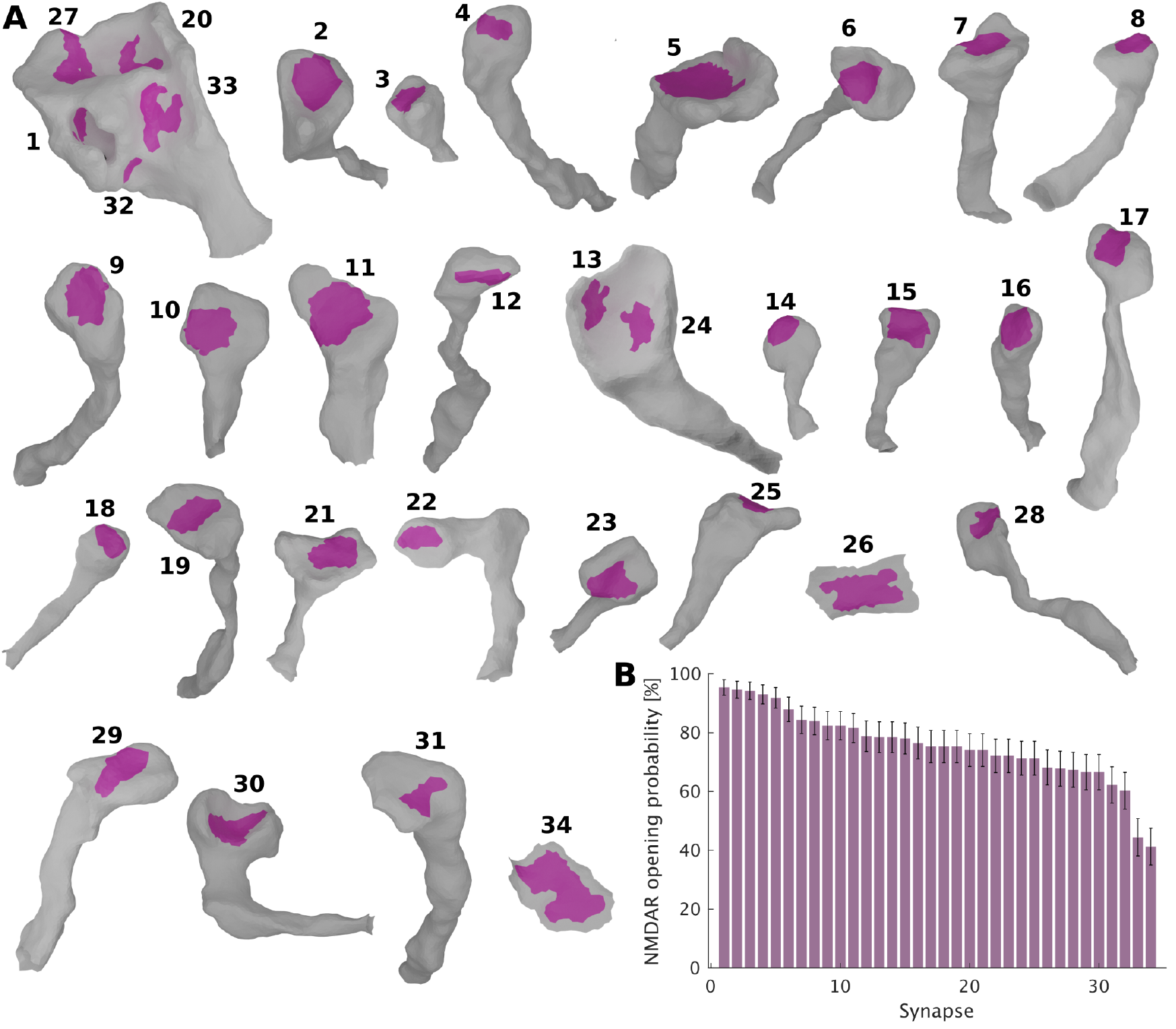
Simulation of NMDAR activation by glutamate assuming realistic synapse geometries reconstructed from mouse cerebral cortex. **A**: This panel displays the 34 reconstructed synaptic geometries utilized in the study, each labeled with a corresponding synaptic number as referenced in panel B. These geometries were derived from dendritic segments imaged by Wu et al. (*32*). **B**: Percentage of simulations in which at least one NMDAR was activated per geometry.

#### 4.1.2. Idealized geometries

To model the idealized geometries, we used spherical caps to represent the curved surfaces and circular meshes for the flat geometries, see schematic in Figure 3A. This approach allowed us to design these geometries with precise control over parameters such as surface area and volume.

**Figure 3:**
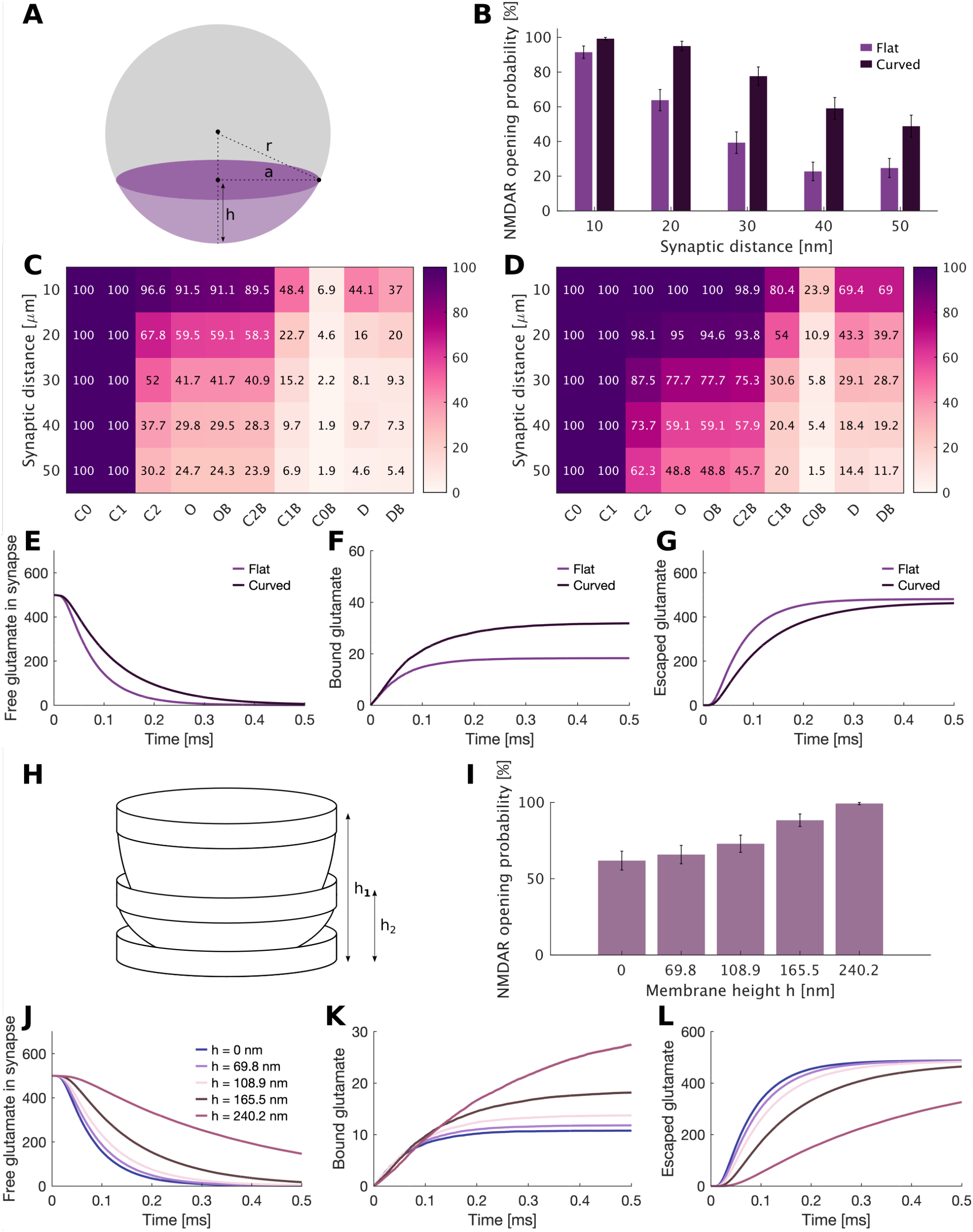
Effect of membrane curvature on glutamate diffusion and NMDAR activation. **A**: Schematic of how the curved meshes are created asuming a spherical cap geometry. **B**: NMDAR opening probability at different synaptic cleft distances for flat and curved membrane geometries. **C**: Percentage of simulations at every NMDAR stage over different synaptic cleft distances for flat membranes. **D**: Percentage of simulations at every NMDAR stage over different synaptic cleft distances for curved membranes. **E**: Mean and standard error of the mean of free glutamate in the synapse for flat and curved membranes. **F**: Mean and standard error of the mean of bound glutamate in the synapse for flat and curved membranes. **G**: Mean and standard error of the mean of escaped glutamate from the synapse for flat and curved membranes. **H**: Schematic of the experiment performed where different curved membranes were considered assuming the same radius a and varying the height h. **I**: NMDAR opening probability assuming different curvatures but a constant radius a and varying height h. **J**: Mean and standard error of the mean of free glutamate in the synapse for different curved membranes. **K**: Mean and standard error of the mean of bound glutamate in the synapse for different curved membranes. **L**: Mean and standard error of the mean of escaped glutamate from the synapse for different curved membranes. SEM is very small and the number of total simulations is n=200.

Although we computed surface area, volume and surface area-to-volume ratio using MCell, the use of idealized geometries enabled us to design these geometries with a targeted surface area-to-volume ratio. To design our curved geometries we build a sphere of radius r, cut it at a height h to obtain a spherical cap of radius a and height h, see schematic in Figure 3A. When controlling for a constant surface area-to-volume ratio across different synaptic cleft distances, we kept the height of the spherical cap h constant and calculated the radius a, see schematic on Figure 3, to achieve the desired surface area-to-volume ratio.

For both realistic and idealized geometries, surface area, volume, and surface area-to-volume ratio were calculated using the Blender *Mesh Analysis* add-on from the MCell4 library (*35*). Our simulations are performed assuming two independent membranes, one for the presynaptic terminal and one for the PSD. However, when referencing the surface area-to-volume ratio of the synapse or calculating the number of glutamate ions within the synapse, we need to consider an enclosed volume. Therefore, we enclose the volume delimited by both membranes and treat this as the synapse. This volume is considered transparent, allowing molecules to diffuse freely in and out. It is used solely for counting the number of glutamate ions within the synapse and calculating the surface area-to-volume ratio of the synapse. See supplementary material for more details.

### 4.2. Signaling model of NMDARs

Throughout this work we assume a constant number of 35 NMDARs on the postsynaptic membrane. This value was chosen to match the density assumed by Bartol et al. (*8*), in the control flat PSDs of Figure 3. We used the NMDAR model proposed by Vargas-Caballero et al. (*33*) and adapted it to dendritic spines as previously done by Bartol et al. (*8*). The equations and parameters used are given in Table 1, and a schematic of the NMDAR model is given in Figure S1B. To activate the NMDARs, both glutamate release and a simultaneous plasma membrane depolarization are necessary. For inducing plasma membrane depolarization, we use the same approach as described in (*7, 8*). The plasma membrane depolarization includes an excitatory postsynaptic potential (EPSP) and a back propagating action potential (BPAP) (*23*), as shown in Figure S1A. For the glutamate release, we assume a point release from the center of the presynaptic membrane (*7*).

**Table 1:**
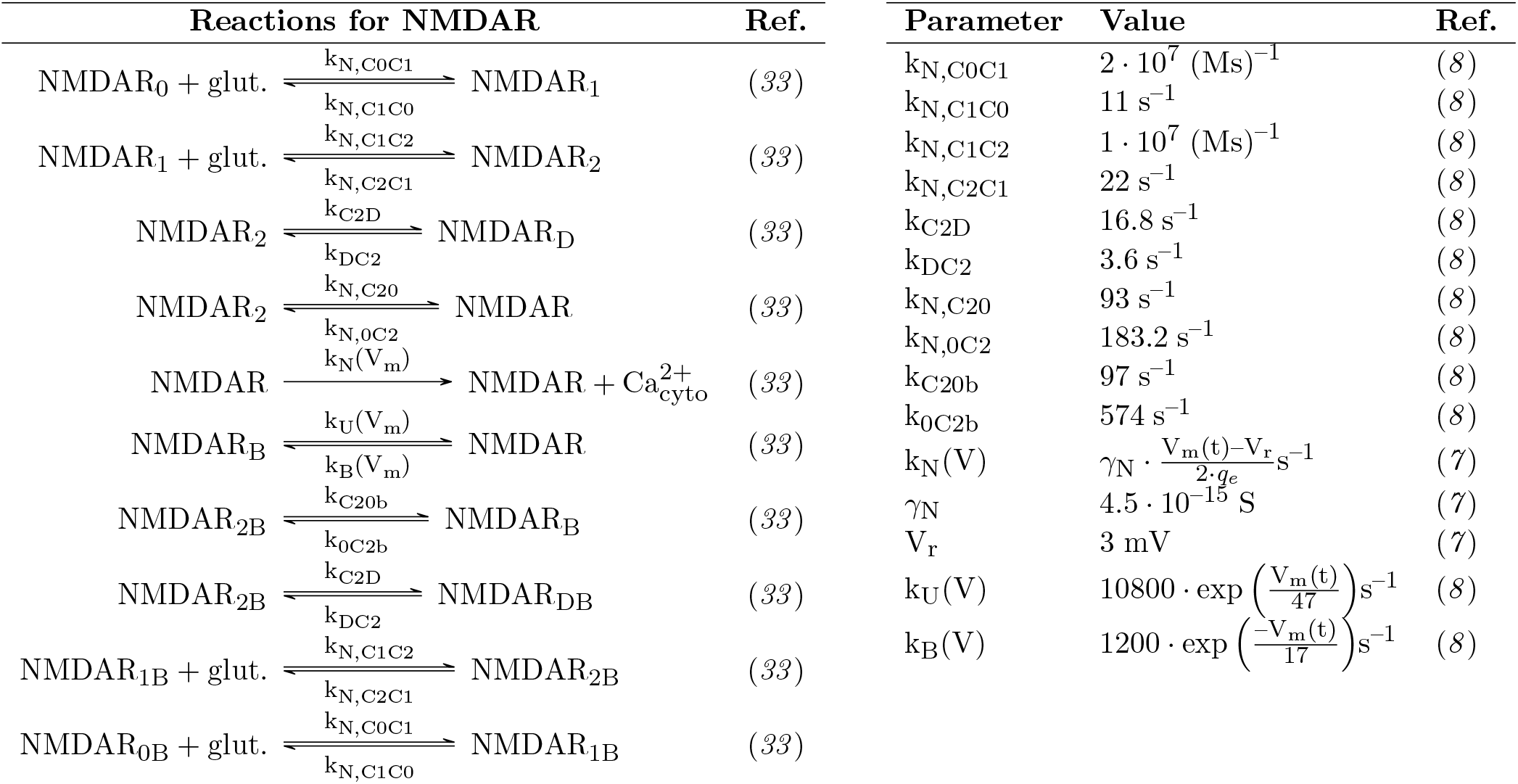
Reactions and parameters used for the NMDAR model.

### 4.3. Metrics Analyzed

During each simulation, we track the number of NMDARs in each model state. We can therefore track how many receptors are open at each timestep for all geometries. We consider the NMDAR opening probability as the number of stochastic simulations (within a set of 250 simulations) containing at least one opened NMDAR.

Within a single simulation, we observed a receptor activation curve containing two peaks, see Figure S1B. For each geometry, we calculated an average curve of open receptors over time and determined both the peak value and the associated standard error of the mean for these curves. This parameter is referre to as the max_NMDAR_ parameter.

The metrics analyzed in this study include several key geometrical and spatial measurements related to synaptic structures and include (a) Synaptic sur face area (SA), (b)Volume (V), (c) Surface area-to-volume ratio (SA/V), (d)Surface area of the postsynaptic density, (SA_PSD_), (e) Average distance from glutamate release to PSD (d_PSD_), (f)Principal curvatures (*κ*_1_ and *κ*_2_) and (g Perimeter (p) and p/SA_PSD_ ratio.

### 4.4. Simulation details

For each simulation setup, we ran 250 simulations in MCell4 (*35*). The glutamate plots, which display the mean and standard error of the mean, were generated using stdshade (*36*) in Matlab version 2022a. The NMDAR opening probability is defined as the percentage of simulations where at least one receptor reaches the open state. For each of the 250 simulations, we compute the number of NMDARs that reach the open state. The maximum number of open NMDARs is calculated by averaging the results from these 250 simulations, identifying the peak of the mean curve, and computing the standard error of the mean at this peak. Both the code and further implementation details will be made publicly available upon publication under github.com/mariahernandezmesa/SynapticCleftCurvature.

## 5 Results

Different geometric features of the synapse can affect synaptic plasticity, including variations in the distance between the presynaptic and postsynaptic membranes, the curvature of the synapse, and the surface area of the PSD (*7, 11, 37, 38*). Here we systematically dissect the role of these parameters in NMDAR activation. We first established the key geometric parameters in our model: the distance between synaptic membranes, membrane curvature, and the surface area of the PSD. To determine the effect of synaptic cleft distance on NMDAR opening probability, we designed a synthetic synapse with two parallel, flat circular membranes, see Figure 1, with different distances. As expected, and in agreemente with previous work from Rusakov et al. (*34*) we observed that an increase in synaptic distance resulted in a decrease in NMDAR opening probability, as illustrated in Figure 1C. Additionally, we noted that increasing the cleft volume led to very small changes in the residence time of glutamate within the synapse, as shown in the inset of Figure 1C. Based on these results from the simplistic model of a synaptic cleft, we next investigated other morphological characteristics affecting particle diffusion and receptor activation, as detailed in the following sections.

### 5.1 Realistic synaptic cleft geometries show significant variability in NMDAR activation

In an effort to uncover the morphological parameters that most influence the activation of receptors within a synaptic cleft, we next focused on realistic reconstructions of dendritic spines. Using the detailed dendritic branch model provided by Lee et al. (*31*), we generated 34 distinct synaptic geometries, shown in Figure 2A. Due to the stochastic nature of MCell simulations, we ran 250 simulations for each geometry to ensure statistical relevance. During each simulation, we track the number of NMDARs in each model state. Figure 2B illustrates the percentage of simulations that resulted in the activation of at least one NMDAR, ranked in descending order. The variance observed was substantial, with activation rates fluctuating between 41.31% and 95.42%. While experimental research has shown that NMDAR activation can lead to morphological changes in synapses (*39*), our results highlight a reciprocally significant influence of synaptic structure on receptor activation dynamics.

In Figure 2B and S2, we analyzed the NMDAR opening probability and max_NMDAR_ and observed that the probability of NMDAR opening is influenced by the shape of the synapse. To further explore the influence of geometry on NMDAR activation, we analyzed a range of geometrical metrics. Specifically, we calculated nine values including the surface area-to-volume ratio and curvature, see Table 2.

**Table 2:** Correlation between different morphological measurements of the realistic meshes and the opening probability and maximum of the mean of the NMDARs. Statistically significant correlations in this plot are indicated as *p *<* 0.05, **p *<* 0.01 and ***p *<* 0.001.

| | SA | V | SA/V | SA<sub>PSD</sub> | d<sub>PSD</sub> | $\kappa_1$ | $\kappa_2$ | p | p/SA<sub>PSD</sub> |
| --- | --- | --- | --- | --- | --- | --- | --- | --- | --- |
| P<sub>open</sub> | -17.46 | *<br>-37.1 | ***<br>58.61 | -15.11 | -33.87 | -1.09 | -7.5 | *<br>-40.63 | -7.41 |
| max<sub>NMDAR</sub> | -9.05 | -29.19 | ***<br>71.81 | -6.88 | -26.88 | 5.91 | -2.99 | -30.73 | -4.66 |

We then assessed the correlation between these parameters and the two above mentioned critical kinetic outcomes related to NMDAR activation: the NMDAR opening probability and the maximum value of the averaged response curve. The results of the correlation study are given in Table 2. Among the geometric parameters examined, only the synaptic volume, the synaptic surface area-to-volume ratio, and the perimeter of the PSD demonstrated a significant correlation with at least one of the parameters quantifying NMDAR activation. Although all three parameters showed correlation with the probability of opening of NMDAR, only the synaptic surface area-to-volume ratio showed a highly significant correlation of 71.81% with the maximum value of the average response curve.

Our results demonstrate a high variability in NMDAR opening probability and indicate that multiple geometric properties likely govern NMDAR opening probabilities in realistic geometries. To systematically explore how different aspects of membrane morphology affect receptor activation, we manipulated idealized geometries to investigate the specific effects of properties such as curvature and surface area-to-volume ratio on glutamate dynamics and NMDAR activation.

### 5.2 Increased synaptic cleft curvature leads to higher NMDAR opening probability

We begin our analysis of idealized geometries of assuming parallel membranes for the presynaptic and postsynaptic membrane and investigating the effects of curvature. Although the analysis of realistic geometries in 2 showed no significant correlation between the curvatures *κ*_1_ and *κ*_2_ and the parameters that quantify NMDAR opening, our previous study indicated that more curved geometries led to greater activation of NMDAR (*23*). This prompted us to investigate how curvature influences NMDAR opening probability in a controlled simulation environment.

A schematic of how the curved meshes were created using the spherical cap geometry is given in Figure 3 A and described in the Methods section. We ran simulations assuming flat and curved parallel membranes, keeping the surface area of both membranes constant. Similar to the approach in Figure 1, we varied the synaptic cleft distance to analyze if the differences between flat and curved synapses persisted across different cleft distances. In Figure 3B, we observed that the NMDAR opening probability was consistently higher in curved synaptic geometries across the range of cleft distances we examined when compared to the flat geometries. To elucidate this result we analyzed the probability of activation for each NMDAR model state, as shown in Figure 3C and D. Across all synaptic distances, there was a significant increase in the likelihood of transitioning to each successive NMDAR state in curved synapses. These results show that curving the synapses increases NMDAR opening probability even when the cleft distance increases when compared to flat membranes. The first significant differences in NMDAR states manifested at the C2 state, with a greater proportion of simulations reaching the C2 state in curved geometries, see Figure 3C and D. Because the transition from C1 to C2 is mediated by glutamate, we hypothesized that curvature of the synapse may enhance the retention of glutamate within the synaptic space. Therefore, we analyzed the glutamate dynamics within the synapse for both morphologies. As shown in Figure 3E, curved synapses were associated with an increased presence of free-diffusing glutamate, which in turn prolonged its residence time. As a consequence, we detected a substantial elevation in the number of glutamate molecules that remained bound to the NMDARs within the curved synapses, coupled with a decrease in the amount of glutamate that escapes the synaptic volume compared to the flat models, as seen in Figures 3F and G, respectively.

Thus far, we have compared curved and flat synapses while maintaining a constant membrane surface area. This constraint resulted in a larger membrane radius, denoted as a, for the flat synapses in contrast to the curved synapses It is well-established that LTP is often accompanied by both an increase in spine size and enhanced exocytosis (*40*). Given this context, we sought to explore the following question: Does augmenting the curvature of a synapse, under the circumstances of increased membrane surface area, correspond to an increased probability of NMDAR activation? To investigate this, we controlled the curvature by adjusting the height of the spherical cap, h, while keeping the radius of the membranes, a, constant. A schematic of how these geometries were built can be seen in Figures 3A and 3H. Our results showed that increasing the curvature led to a higher probability of NMDAR opening, as previously observed, see Figure 3I. Furthermore, we observed an increase in the levels of free diffusing glutamate and bound glutamate, as well as a decrease in the amount of glutamate that escaped the synaptic space, see Figures 3J, 3K and 3L. These findings highlight the conclusion that synaptic curvature does not only influence receptor activation, but can also impact neurotransmitter dynamics, ultimately contributing to the synaptic activation in response to increased cleft distances.

### 5.3 Maintaining a constant surface area-to-volume ratio altered NMDAR opening probability for flat geometries

In Figure 3, we explored how curvature impacts NMDAR kinetics and glutamate diffusion, noting that increased curvature also raises the synapse surface area-to-volume ratio. As we observed in our analysis of realistic synapse geometries (see Table 2), the surface area-to-volume ratio of the synapses can affect NMDAR opening and glutamate diffusion. To isolate these effects, we simulated two flat membranes with varying cleft distances, while holding a constant surface area-to-volume ratio of 102 *μ*m by adjusting membrane radii (Figure 4A).

**Figure 4:**
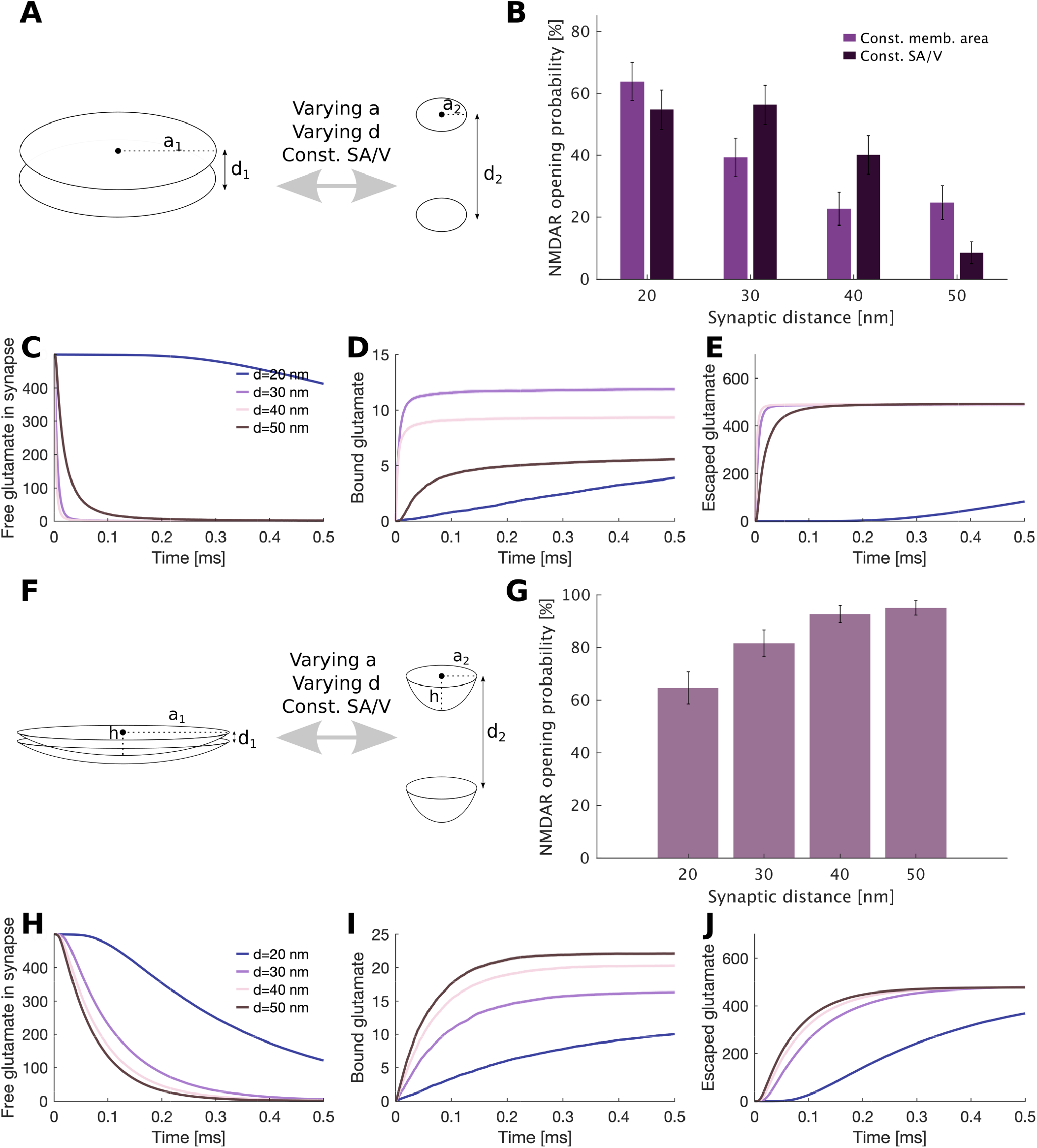
NMDAR opening probability and glutamate diffusion assuming a constant surface area-to-volume ratio at different synaptic cleft distances. **A**: Assuming flat membranes we varied the radius a to ensure constant surface area-to-volume ratio at different synaptic cleft distances. **B**: The NMDAR opening probabilities for flat membranes assuming a constant membrane area as in Figure 1 and varying the radius a to ensure a constant surface area-to-volume ratio as shown in A. **C**: Mean and standard error of the mean of free glutamate in the synapse for flat and curved membranes. **D**: Mean and standard error of the mean of bound glutamate in the synapse for flat and curved membranes. **E**: Mean and standard error of the mean of escaped glutamate from the synapse for flat and curved membranes. **F**: Assuming curved membranes we varied the radius a, keeping the height of the spherical cap h constant, to ensure constant surface area-to-volume ratio at different synaptic cleft distances. **G**: The NMDAR opening probabilities for curved membranes when varying the radius a to ensure a constant surface area-to-volume ratio as shown in F. **H**: Mean and standard error of the mean of free glutamate in the synapse for flat and curved membranes. **I**: Mean and standard error of the mean of bound glutamate in the synapse for flat and curved membranes. **J**: Mean and standard error of the mean of escaped glutamate from the synapse for flat and curved membranes.

In comparison to the geometries with a constant surface area presented in Figure 1, the surface area-to-volume ratio was higher for synaptic cleft distances ranging from 30 – 50 nm in this experiment, but lower at 20 nm. NMDAR opening probabilities are shown in Figure 4B. It is worth noting that assuming a constant surface area-to-volume ratio resulted in higher NMDAR opening probabilities at synaptic cleft distances of 30 nm and 40 nm. However, at 20 nm, the NMDAR opening probability was slightly, though not significantly, smaller compared to Figure 1. This discrepancy could be attributed to both the lower surface area-to-volume ratio and to the larger membrane surface area, resulting in a longer residence of glutamate within the synapse, as shown in Figure 4C. However, despite the longer residence time within the synapse, it is important to note that the binding rate of glutamate was lower, as shown in Figure 4D. This observation can be attributed to the fact that glutamate must diffuse over longer distances to activate the NMDARs.

Although maintaining a constant surface area-to-volume ratio helped compensate for the effects of synaptic distances at short ranges between 30 nm and 40 nm, it was unable to completely compensate for the inverse relationship between cleft distance and NMDAR opening probability, see 50 nm in 4B. As a result, an overall decrease in the opening probability was observed at larger cleft distances. Furthermore, at 50 nm, we noticed a decrease in the NMDAR opening probability compared to the simulations shown in Figure 1.

### 5.4 Increased curvature in synaptic membranes mitigates the effects of synaptic distance when constant surface area-to-volume ratio is maintained

We next aimed to explore the interplay between membrane curvature and the surface area-to-volume ratio. Building on the experiment shown in Figures 4A-E, we maintained a constant surface area-to-volume ratio in curved geometries while varying synaptic distances d_i_.To achieve this, we kept the height *h* of the spherical cap geometry constant and adjusted its radius a accordingly, as illustrated in Figure 4F. This approach results in increased membrane curvature at greater synaptic cleft distances.

The results of our simulations show that at greater synaptic cleft distances, NMDAR opening probability increases. These findings contrast with those from Figure 3B, where, under conditions of constant curvature and varying surface area-to-volume ratios, the NMDAR opening probability decreased with increasing synaptic cleft distance. Thus, our results indicate that increased curvature can compensate for the decrease in NMDAR opening probability due to increased synaptic cleft distance. Interestingly, as observed in Figures 4C-E, despite longer glutamate residence time at shorter synaptic cleft distances (see Figure 4H), the amount of bound glutamate was lower compared to that at greater synaptic cleft distances. We hypothesize that this may be due to an increased membrane surface area at lower synaptic cleft distances. However, designing simulations that maintain constant surface area, surface area-to-volume ratio, and curvature simultaneously is not feasible. Nevertheless, our study reveals that with a constant surface area-to-volume ratio, increased curvature leads to a higher NMDAR opening probability.

### 5.5 Presynaptic and postsynaptic membrane morphology determine the surface area-to-volume ratio of the synaptic cleft and thereby affect NMDAR opening probability

Up to this point, our simulations have assumed identical, parallel membranes for both the presynaptic and postsynaptic components. However, we wanted to understand the impact of non-parallel, heterogeneous membranes on NMDAR activation and glutamate diffusion. Therefore, we ran four sets of simulations assuming concave and convex presynaptic membranes, as well as concave and convex PSDs paired with flat presynaptic membranes. These configurations are shown in Figure 5.

**Figure 5:**
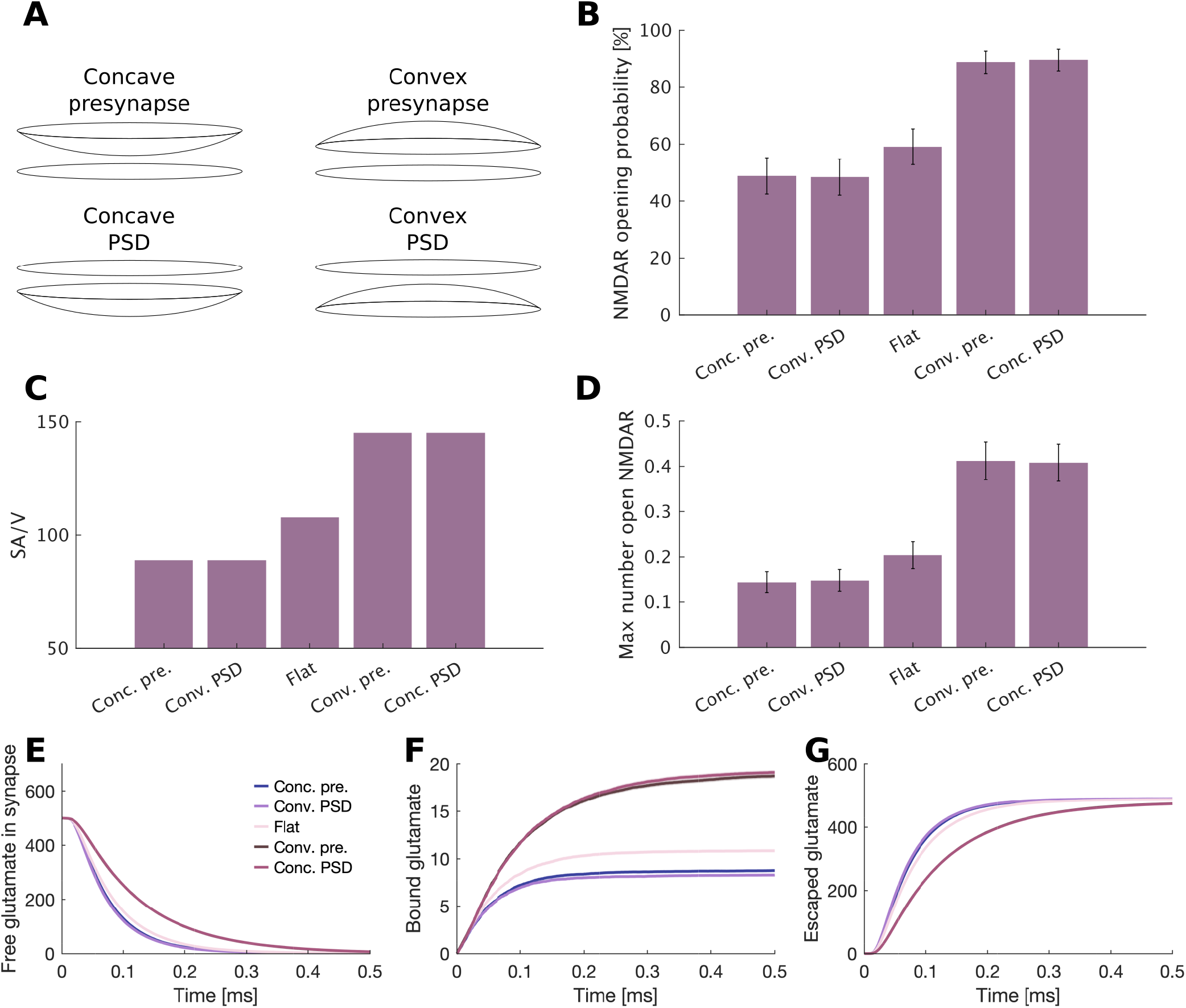
Simulations assuming non parallel presynaptic and postsynaptic membranes. **A**: Schematic showing the different geometries used for the experiment. **B**: The NMDAR opening probabilities for the geometries assuming non parallel membranes for the presynaptic membrane and the PSD. **C**: Surface area to volume raio of the different geometries. **D**: Maximum of the mean of the opened NMDARs calculated over time. **E**: Mean and standard error of the mean of free glutamate in the synapse. **F**: Mean and standard error of the mean of bound glutamate in the synapse. **G**: Mean and standard error of the mean of escaped glutamate from the synapse.

We observed that the highest NMDAR opening probability occurs when assuming the convex presynapse or the concave PSD, see Figure 5B. Conversely, the lowest probability of NMDAR opening likelihood is associated with a concave presynapse or a convex PSD configuration. Identical results were obtained when analyzing the maximum number of opened NMDARs, see Figure 5D. These results correspond to the varying surface area-to-volume ratios of these geometries, as shown in Figure 5C, where a greater ratio enhances NMDAR activation efficiency.

The increased surface area-to-volume ratio translates to an increased synaptic residence time for free glutamate in the convex presynapse and concave PSD experiments (see Figure 5E). This prolonged glutamatergic residence time increases the probability of glutamate binding, which in turn leads to a higher accumulation of bound glutamate (see Figure 5G) and simultaneously limits the leakage of glutamate from the synaptic cleft (see Figure 5G).

From these findings, it becomes apparent that the influence on synaptic communucation efficiency stems not just from the differences between the presynaptic and postsynaptic membranes in isolation, but also from the collective synaptic volume and morphology of the synaptic space. Confirming this, simulations applying parallel concave and parallel convex membrane arrangements yielded congruent results, see Figure S4.

### 5.6 NMDAR clustering increases receptor activation and is more robust against changes in synaptic morphology

NMDARs are known to aggregate within the PSD of dendritic spines (*41*). To explore the impact of NMDAR clustering on the synaptic morphology effects on NMDAR opening probability, we revised certain geometries previously examined but with a new constraint including NMDAR clustering. This was simulated by localizing NMDARs to a specific confined area of the PSD, demarcated by the purple region in Figure 6. We replicated the following experiments assuming clustering: (a) we varied the membrane curvature while keeping the radius (a) constant, recapitulating conditions described in Figure 3, (b) we modified the radius (a) to preserve a constant surface area-to-volume ratio, following the protocol outlined in Figure 4, and (c) we replicated the asymmetric or non-parallel configurations as initially presented in Figure 5. In all cases, we observed an increase in NMDAR opening probability when clustering the NMDARs, see Figure 6. Our approach allowed us to methodically assess how the physical congregation of NMDARs within the PSD influences synaptic transmission under diverse geometric conditions. Our results show that clustering the NMDARs towards the release location of glutamate from the presynaptic membrane not only increases NMDAR opening probability, but also reduces the effects of synaptic morphology on NMDAR activation. Furthermore, clustering the NMDARs allows glutamate unbinding from the NMDARs to bind to neighboring receptors and thereby induce their activation, enhancing receptor cooperation.

**Figure 6:**
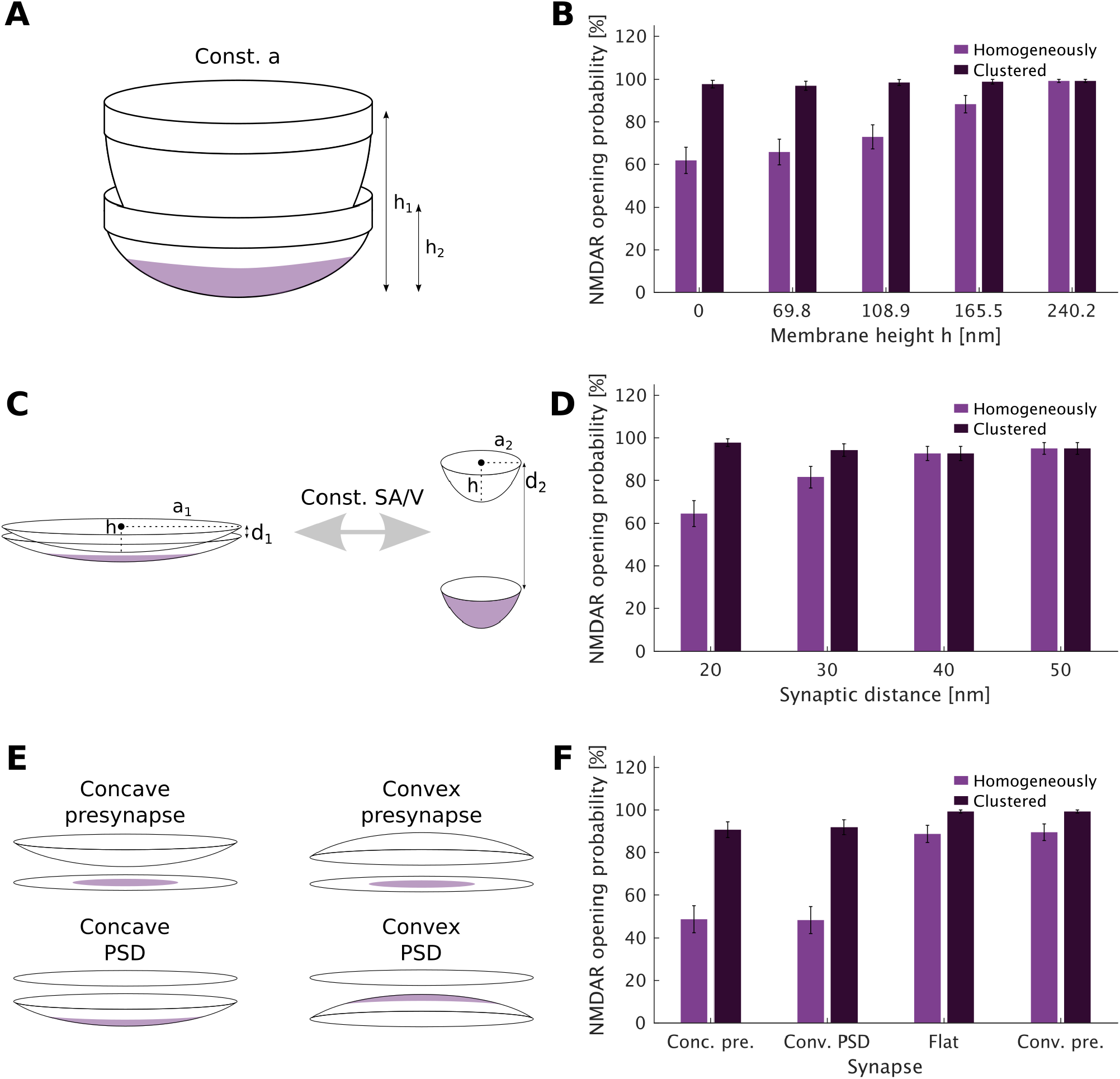
Homogeneously distributed versus clustered NMDAR distribution in different synaptic geometries. **A**: Schematic showing clustering when assuming different curvatures, by keeping the radius a constant and varying the height h of the spherical cap. **B**: NMDAR opening probability for the geometries of A comparing homogeneously distribution of the NMDARs and clustered distribution. **C**: Schematic showing clustering when controlling for a constant surface area-to-volume ratio, by varying the radius a and keeping the height h of the spherical cap constant. **D**: NMDAR opening probability for the geometries of C comparing homogeneously distribution of the NMDARs and clustered distribution. **E**: Schematic showing clustering when assuming non parallel membranes for the presynaptic membrane and the PSD. **F**: NMDAR opening probability for the geometries of E comparing homogeneously distribution of the NMDARs and clustered distribution.

## 6 Discussion

In this study, our aim was to investigate the influence of dendritic spine morphology on glutamate diffusion and subsequent activation of NMDARs. Addressing this challenge involves studying the complexities presented by variations in membrane curvature, morphology, surface area, and the volume contained by synaptic membranes. Our findings demonstrate that the curvature of the membrane and the surface area-to-volume ratio of synaptic junctions significantly modulate the efficiency of synaptic transmission. Diverse geometrical models, both idealized and reconstructed from EM images, revealed marked disparities in receptor activation patterns, underscoring the critical role of spatial configuration in synaptic function. Furthermore, we demonstrate that NMDAR clustering not only increases NMDAR activation, in alignment with previous studies (*42, 43*), but also mitigates the effects of synaptic shape on receptor activation.

Our study shares conceptual roots with the seminal work by Rusakov et al. (*34*) on extrasynaptic glutamate diffusion in the synaptic cleft, which modeled diffusion with a focus on ultrastructural constraints. Our findings on the effect of increased glutamatergic release distance reducing NMDAR activation align with those reported by Rusakov et al.(*34*). However, key methodological differences distinguish our approach. Rusakov’s work utilized a deterministic framework to model diffusion, while our simulations employ a particle-based, stochastic approach. By avoiding a deterministic approach, our study provides a closer approximation of glutamate diffusion dynamics within the highly variable synaptic microenvironment, where particle-based models can better account for the probabilistic nature of neurotransmitter-receptor interactions. Moreover, unlike Rusakov’s explicit focus on the extracellular space’s porosity and tortuosity, our model does not explicitly simulate these properties (*34*). Our modeling approach includes bulk surface coupling boundary conditions to represent interactions at both the presynaptic release site and the postsynaptic membrane, as well as diffusive flux at adjacent boundaries. This detailed consideration of boundary interactions within the synaptic cleft provides a unique insight into glutamate diffusion and receptor activation.

The diminutive size of the synaptic cleft, coupled with the increased number of neurotransmitter particles released during synaptic activity, poses a significant challenge for imaging these structures. Stochastic spatial simulations provide a powerful approach to explore how synaptic geometry, as defined by the boundaries of the presynaptic and postsynaptic membranes, influences patterns of receptor activation. Although our focus here is on synaptic applications, we believe that understanding how the shape of a small volume delimited by two membranes affects receptor activation can also be interesting for other biological problems. Here, we have also highlighted the significant relation between synaptic morphology and receptor activation. Given that the physical structure of the synaptic cleft is a key determinant in the process of signal transduction, modelers should consider including multiple synaptic geometries when drawing conclusions from their biophysical simulations. The implications of this study also suggest that biological experimentation should expand its scope to include an assortment of synaptic geometries. This is particularly relevant since the diffusion trajectories and the retention time of molecules like glutamate within synaptic spaces appear to be significantly influenced by the geometry of the surrounding membranes.

While the current study provides significant insights into the interplay between membrane morphology and signal transduction, it also illuminates several aspects that should be considered for future work. Firstly, our findings demonstrate a considerable correlation between membrane curvature and NMDAR opening probability within idealized synaptic models. However, realistic geometries reconstructed from experimental images are much more complex including diverse curvature distributions as outlined in previous works (*23, 44*). An analysis of the average principal curvatures, *κ*_1_ and *κ*_2_, within these complex architectures revealed no discernible link to NMDAR opening dynamics (see Figure 2). Further analysis of the maximum value of the principle curvatures within this complex geometries revealed no significant correlation. We believe that the intrinsic complexity of realistic synaptic geometries renders simple metrics such as average or maximum curvature values insufficient for capturing their influence on receptor behavior. Furthermore, the realistic geometries are very complex with varying curvature, shape and surface area of the membranes. Future studies should address this concern and try to understand the relation between curvature and receptor activation in realistic geometries.

Secondly, NMDAR activation requires both glutamate binding and plasma membrane depolarization. In our computational model, we controlled for the latter by assuming constant depolarization across various morphologies. However, we must not overlook the potential impact of curvature on electrodiffusive processes (*45*). Therefore, future work should consider combining a stochastic reaction diffusion framework with an electrodiffusion model to better comprehend the influence of membrane topography on signal propagation.

Recently, Li et al. (*46*) used Monte Carlo simulations and mean field theory to explore the dynamics of protein condensation and clustering on cell membranes, specifically alongside the membranes undulatory motions. They show how alterations in the height and width of periodic membrane topography can influence the agglomeration of proteins within the membrane (*46*). Building on the findings of Li et al. (*46*), we propose that integrating their simulation approach into our studies of particle diffusion and receptor activation within small volumes limited by membranes may yield deeper insights. This could significantly advance our understanding of how membrane morphology not only modulates particle diffusion within the confined volume but also affects receptor clustering and the subsequent triggering of receptor activation.

In addition to NMDAR activation, glutamate diffusion and its uptake by astrocytes play critical roles in synaptic function. Given the proximity of astrocytic processes to synaptic sites, these cells contribute to maintaining glutamate homeostasis by uptaking excess glutamate, thereby shaping synaptic signaling. Our results emphasize how spatial constraints and geometry influence glutamate diffusion and receptor activation, a relationship that is likely modulated *in vivo* by astrocytic uptake dynamics. Future studies integrating astrocytic glutamate uptake in conjunction with synaptic morphology may further elucidate the nuances of synaptic transmission, especially under varying physiological and pathological conditions.

We summarize by noting that several biological processes share similarities with glutamate release into the synaptic cleft. Thus, in addition to delineating the relationship between dendritic spine morphology and glutamate diffusion and NMDAR activation, our work can serve as motivation to study the interaction of membrane shape and signaling efficiency in similar biological contexts. For example, healthy cardiomyocytes contain curved plasma membrane invaginations (T tubules), allowing proximity between L-type Ca^2+^ channels (LTCCs) in the plasma membrane and ryanodine receptors (RyRs) on the sarcoplasmic reticulum (SR). The confined space between the plasma membrane and SR, termed the dyadic cleft, optimizes Ca^2+^ signaling. The proximity between the LTCCs and the RyRs in the dyadic cleft allows for an enhanced release of Ca^2+^ from the ER, a process crucial to the excitation-contraction coupling mechanism of the heart. The shape of the T-tubules, including its curvature, is known to be disrupted during heart failure, a factor that can perturb Ca^2+^ dynamics (*47*). The alterations in the geometry of the cardiac dyad, much like changes in synaptic geometry, could have profound effects on Ca^2+^ diffusion and triggering of RyRs, drawing a parallel to the impact of synaptic structure on glutamate diffusion and NMDAR activation. Further examples of biological processes where the confined space between two membranes could influence receptor activation include the neuromuscular junctions, immune synapses and the intermembrane space of mitochondria. Thus, detailed spatial models can help us uncover shared biophysical principles across different biological systems.

## 7 Acknowledgments

MHM and KJM are supported by the Simula-UCSD-University of Oslo Research and PhD training (SUURPh) program, an international collaboration in computational biology and medicine funded by the Norwegian Ministry of Education and Research. This work was supported in part by Air Force Office of Scientific Research (AFOSR) MURI FA9550-18-1-0051 to PR.

## 8 Supplemental material

### 8.1 Calcualations for the geometries based on the sperical cap equations

The surface area of the spherical cap is given by

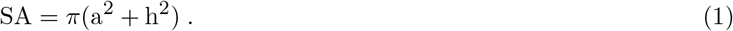

The idealized curved geometries consist of parallel spherical caps for both the presynaptic membrane and the PSD. Therefore, the surface area of the enclosed synaptic volume is calculated as twice the surface area of the spherical cap, plus the walls of the synapse, which are calculated assuming cylindrical geometries

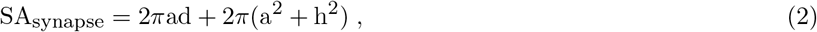

where d is the height of the synaptic cleft, defined as the distance between the presynaptic membrane and the PSD.

The volume of the synapse is defined as

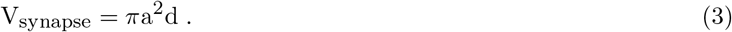

The surface area-to-volume ratio is calculated by dividing equation 2 by equation 3. Knowing that the relationship between the spherical radius r, the radius of the spherical cap a and the height of the spherical cap h is

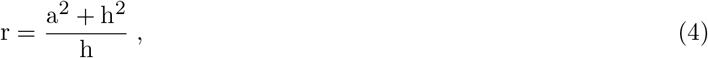

we can, for example, assume a constant height h and calculate the radius of the spherical cap a to obtain a desired surface area-to-volume ratio for the synaptic volume.

**Figure S1:**
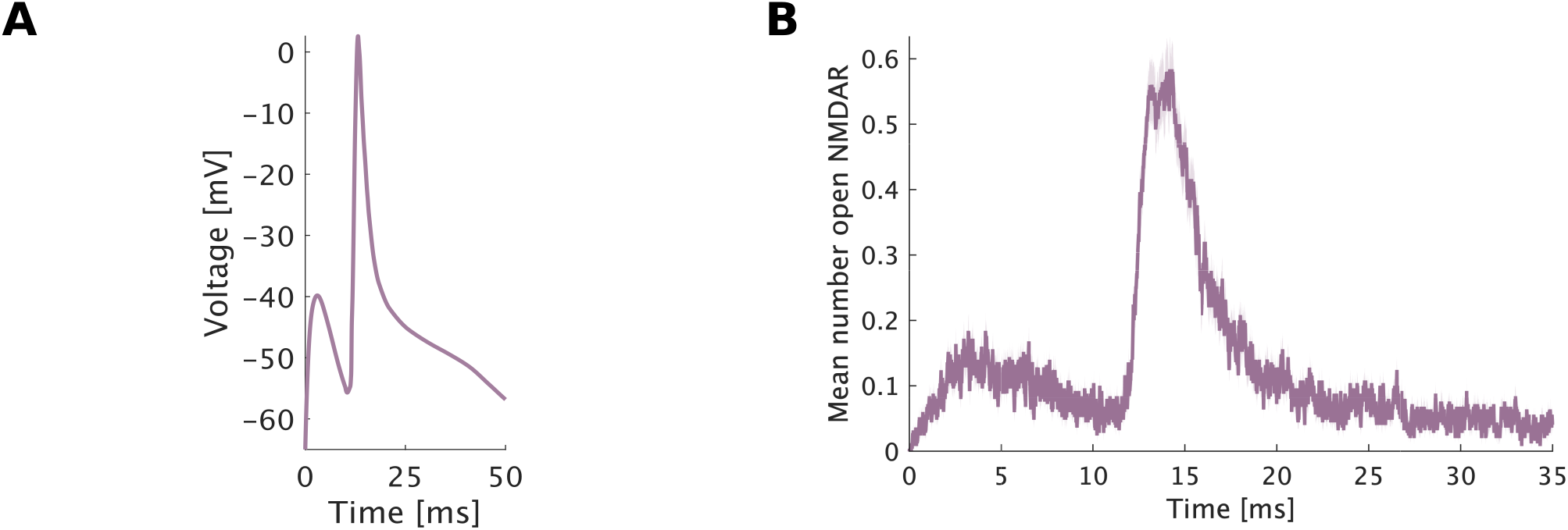
Schematic of the NMDAR model and average of open NMDAR over 250 simulations. **A**: Schematic of NMDAR reaction model proposed by (*33*). **B**: An example of the mean and standard error of the man curve of the number of opened NMDARs over 250 simulations.

**Figure S2:**
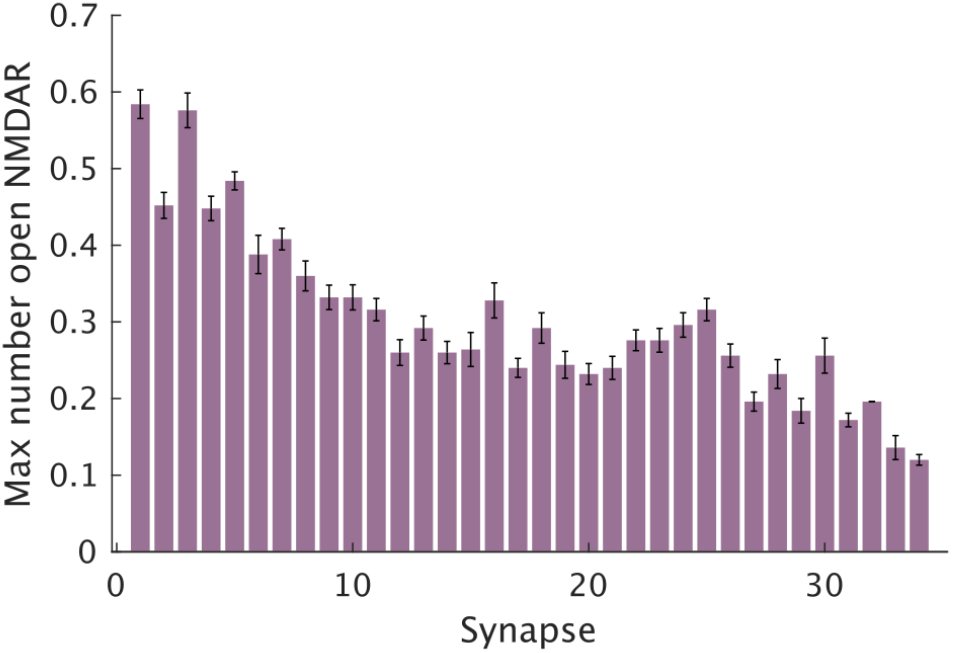
Maximum of the mean of the opened NMDARs calculated over time.

**Figure S3:**
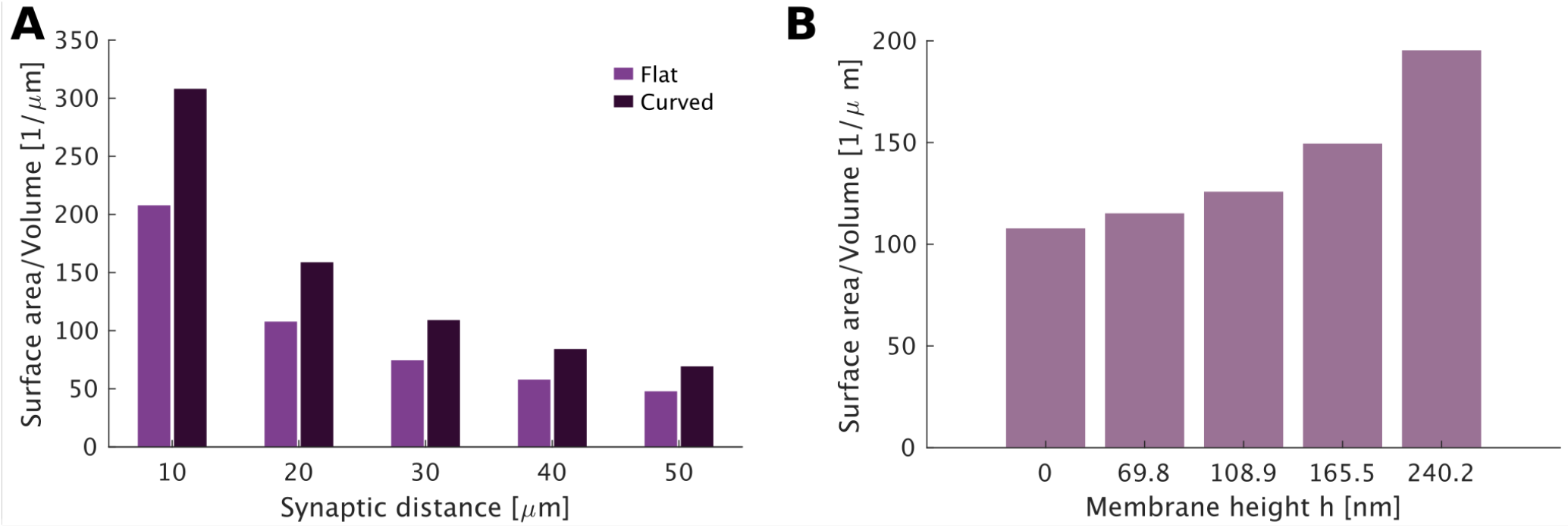
Surface area-to-volume ratio of the geometries in Figure 3. **A**: Suface area to volume ratio of the geometries used in the simulations for Figure 3B. **B**: Suface area to volume ratio of the geometries used in the simulations for Figure 3I.

**Figure S4:**
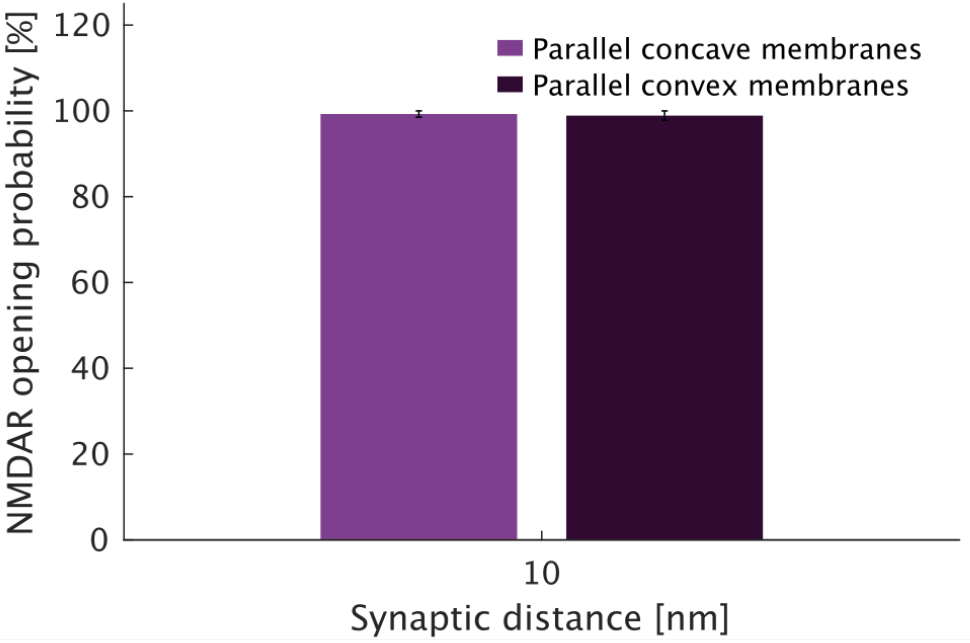
Comparison between parallel concave and parallel convex membranes.

